# Representing Genetic Determinants in Bacterial GWAS with Compacted De Bruijn Graphs

**DOI:** 10.1101/113563

**Authors:** Magali Jaillard, Maud Tournoud, Leandro Lima, Vincent Lacroix, Jean-Baptiste Veyrieras, Laurent Jacob

## Abstract

**Motivation:** Antimicrobial resistance has become a major worldwide public health concern, calling for a better characterization of existing and novel resistance mechanisms. GWAS methods applied to bacterial genomes have shown encouraging results for new genetic marker discovery. Most existing approaches either look at SNPs obtained by sequence alignment or consider sets of kmers, whose presence in the genome is associated with the phenotype of interest. While the former approach can only be performed when genomes are similar enough for an alignment to make sense, the latter can lead to redundant descriptions and to results which are hard to interpret.

**Results:** We propose an alignment-free GWAS method detecting haplotypes of variable length associated to resistance, using compacted De Bruijn graphs. Our representation is flexible enough to deal with very plastic genomes subject to gene transfers while drastically reducing the number of features to explore compared to kmers, without loss of information. It accomodates polymorphisms in core genes, accessory genes and noncoding regions. Using our representation in a GWAS leads to the selection of a small number of entities which are easier to visualize and interpret than fixed-length kmers. We illustrate the benefit of our approach by describing known as well as potential novel determinants of antimicrobial resistance in *P. aeruginosa,* a pathogenic bacteria with a highly plastic genome.

**Availability and implementation:** The code and data used in the experiments will be made available upon acceptance of this manuscript.

## 1 Introduction

Antimicrobial resistance has become a major worldwide public health concern, as illustrated by the increase of hospital-acquired infections on which both empirical and targeted treatments fail because of multi-resistant bacterial strains (Micek *et al*., 2015). This worrisome situation calls for a better comprehension of the genetic bases of resistance mechanisms. Genome-wide association studies (GWAS) aim at linking genetic determinants to phenotypes, and seem appropriate for this purpose. Indeed over the past four years, bacterial GWAS have shown encouraging results for genetic marker discovery thanks to the increase in rich panels of bacterial genomes and phenotypic data (Alam *et al*., 2014; Chewapreecha *et al*., 2014; Earle *et al*., 2016; Farhat *et al*., 2013; Sheppard *et al*., 2013).

GWAS rely on a particular definition of genetic variants, such as the presence in the genome of SNPs against a reference genome, of genes in a predefined list or of fixed-length kmers. Each genome in the panel is encoded as a vector with one entry per genetic variant - indicating, *e.g*., whether the genome contains the variant - and all variants are tested for association with the phenotype of interest. The objective of this paper is to describe a novel representation of genetic variation for bacterial GWAS, and to discuss its advantages over existing ones.

Most existing bacterial association studies use approaches developed for human GWAS to encode genome variation: they align all genomes in the panel against a reference genome, identify SNPs and represent each strain by a presence/absence vector with one entry per SNP (Farhat *et al*., 2013; Alam *et al*., 2014; Chewapreecha *et al*., 2014). However a suitable reference is not always available, in particular for species with extensive genome plasticity and a large accessory genome. The accessory genome is the part of the genome not found in all strains of the same species, and is largely composed of genetic material acquired by horizontal gene transfer. For highly plastic species – including pathogenic and antibiotic resistant species such as *P. aeruginosa–,* it can represent more than a quarter of the complete genome, leading to manifold genomes which vary by their size and content (Kung *et al*., 2010). Aligning such genomes against a reference makes little sense and alternative representations of genetic variation are required.

To account for the variation in gene content, some studies also use as candidates the presence or absence of genes represented in the studied panel (Earle *et al*., 2016). However, genetic determinants linked to transcriptional or translational regulation may be located in noncoding regions, and thus are missed by approaches relying on this representation, whose quality also depends a lot on the quality of the available annotation.

Finally to get around these issues, other studies have represented genomes as vectors of presence or frequency of kmers, *i.e.,* of length *k* sequences in the genome (Sheppard *et al*., 2013; Earle *et al*., 2016). Contrarily to SNP- or gene-based approaches, kmers are able to describe genome diversity without requiring an alignment against a reference genome or prior annotation. A major issue of this approach however is that the number of distinct kmers contained in a set of genomes increases with k, easily reaching tens of millions of candidates and leading to very high memory requirements, time loads and complexity in feature interpretation. At the same time it is clear that the information carried by kmers is highly redundant as each single locus is represented by several overlapping kmers, suggesting that they are not the optimal resolution to describe genome variation. Thus, the best way to encode genomic variation in bacterial GWAS remains an open question (Read and Massey, 2014; Power *et al*., 2017).

Our proposed representation is based on compacted De Bruijn graphs (de Bruijn, 1946) (DBG), which are widely used for *de novo* genome assembly (Pevzner *et al*., 2001; Zhang *et al*., 2011) and variant calling (Iqbal *et al*., 2012; Le Bras *et al*., 2016). All fixed-length kmers corresponding to the same long sequence in a set of genomes are represented as a single longer word associated with a node in the graph. The nodes of the compacted DBG therefore provide a lossless, data dependent compression of the fixed-length kmers, leading to a resolution adapted to the local variability of the genomes.

We show in this paper how using these nodes to define genetic variants for bacterial GWAS indeed leads to selecting a few entities which are easier to interpret and make more sense biologically than fixed-length kmers. We also show how DBGs themselves facilitate the analysis of a set of candidate variants found to be significantly associated with microbial antibiotic resistance. We illustrate these advantages using a panel of *P. aeruginosa* strains with multiple phenotypic resistances to antimicrobial drugs.

## 2 Methods

We here introduce our proposed definition of genomic variants, showing how it generalizes two standard alternatives based on presence/absence of:

- SNPs obtained by alignment against a reference genome,
- Fixed-length kmers.

We then detail how we use it in a GWAS context and how we assess its performance.

### 2.1 Encoding genome variation using compacted De Bruijn graphs

DBGs are directed graphs representing overlaps between a set of strings. More specifically, the DBG nodes are all unique kmers extracted from the sequences and an edge is drawn between any two nodes if the (*k*-1)-length suffix of one equals the (*k*-1)-prefix of the other.

When considering a set of similar sequences, a single DBG built over all these sequences displays a particular topology, providing information on any variation among sequences in the set. A SNP for example leads to kmers which are constant across genomes, followed by kmers differing by one letter, followed by more constant kmers. When building the DBG, if a kmer overlaps two other kmers but these two kmers differ by their *k*th base we obtain a fork pattern in the graph (Figure 1A). When both branches of the fork join again into one shared kmer, we obtain a bubble pattern with two branches of equal length, representing the SNP (Figure 1B). Insertions of large sequences in some of the genomes lead to bubbles with one branch longer than the other, and can therefore be represented in the same framework. This makes DBGs a tool of choice to describe genomic variants (Le Bras *et al*., 2016).

**Figure 1:**
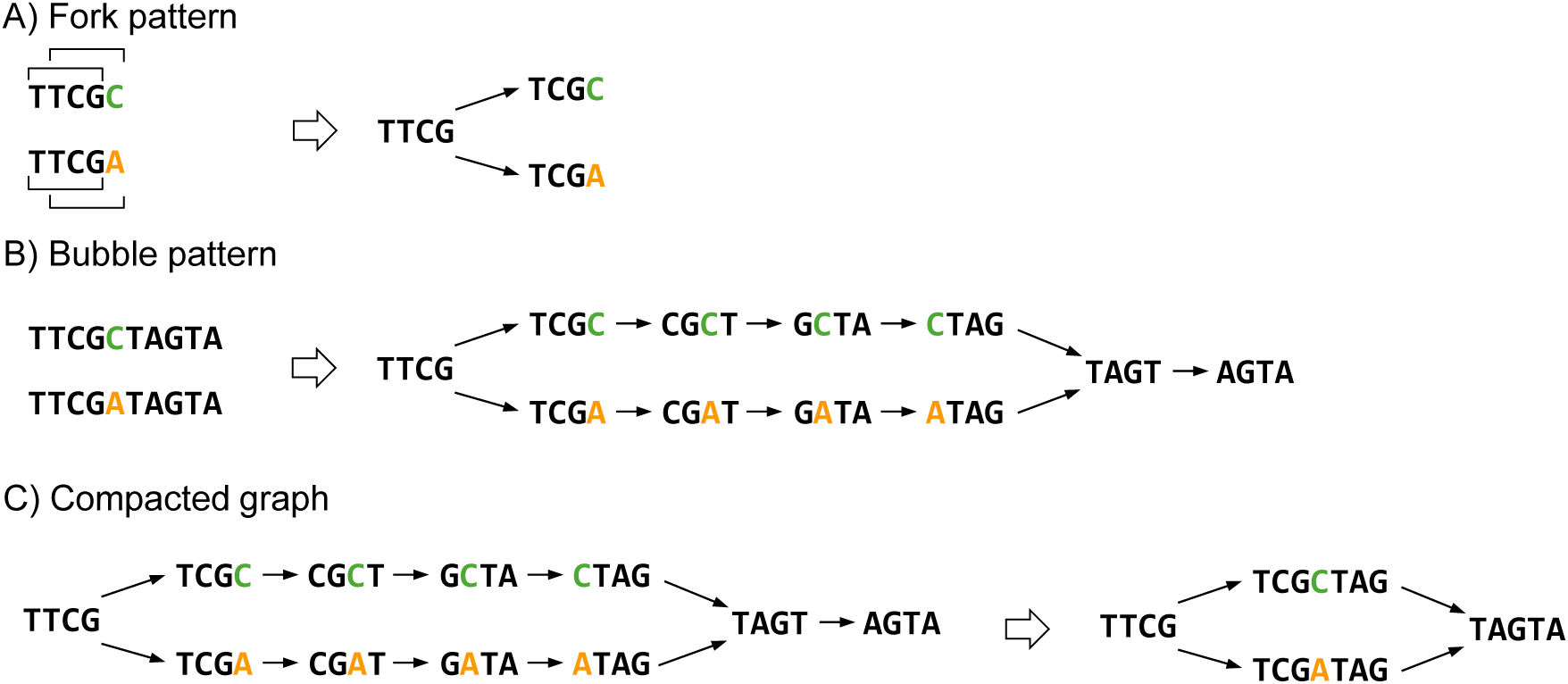
DBG construction. For this example, k=4. A) the 4-mer “TTCG” present in both sequences overlaps two other 4-mers (“TCGC” and “TCGA”) but these two 4-mers differ by their 4th base and we obtain a fork pattern. B) both branches of the fork join on the shared 4-mer “TAGT”, and this creates a bubble pattern representing here the SNP C to A. C) linear paths of the graph are compacted; the remaining graph contains fewer nodes representing longer kmers (unitigs): two 4-mers and two 7-mers instead of eleven 4-mers before compaction. Compacted nodes have variable length.

Interestingly, these graphs can be compacted by first using a unique node to store a kmer sequence and its reverse complement, and then merging linear paths, *i.e.,* sequences of nodes not linked to more than two other nodes. This compression is done without loss of information, because it only affects redundant descriptors, *i.e*., kmers whose presence/absence pattern is identical across genomes (Butler *et al*., 2008; Zerbino and Birney, 2008; Chikhi *et al*., 2016). Thus, the nodes of the compacted DBG can be thought of as haplotypes of variable length in different regions of the genomes, including coding and noncoding regions as well as core and accessory genome (Figure 1C). In the remainder, we denote by unitig the variable-length kmer associated with a node in the compacted graph.

Rather than representing genomes by presence/absence patterns of SNPs, full genes or fixed-length kmers, we propose to use presence/absence patterns of these unitigs. We discuss in Section 2.1.1 how they generalize in an adaptive fashion existing representations based on presence/absence patterns of fixed length kmers or of SNPs defined by alignments against a reference genome.

#### 2.1.1 Unitigs, SNPs and fixed-length kmers

When dealing with a clonal panel of very similar genomes, genomic variants in prokaryote genomes are classically defined as the presence/absence of SNPs identified by alignment of each genome against a reference. For highly plastic genomes on the other hand, alignment against a single reference genome is unsuitable and genomic variation is often encoded as the presence/absence of fixed-length kmers in the genomes. The presence/absence of unitigs of a DBG built over the genomes of several individuals provides a flexible representation thereof which interpolates between these two alternatives in a data adaptive fashion.

At one extreme in the case of a clonal panel with only SNPs as genetic variants (Figure 2A), the DBG is a path with a few bubble patterns - assuming genomes do not contain repeated regions longer than k. This graph is isomorph to a reference genome with SNPs. On the other hand, the collection of fixed-length kmers belonging to these genomes is very redundant: all kmers containing the same SNP at different positions have the same presence/absence across strains by construction. Those containing no SNP - most of them - do not represent any polymorphism: they are present in all genomes in the panel and their presence/absence representation would be 1 identically across strains.

**Figure 2:**
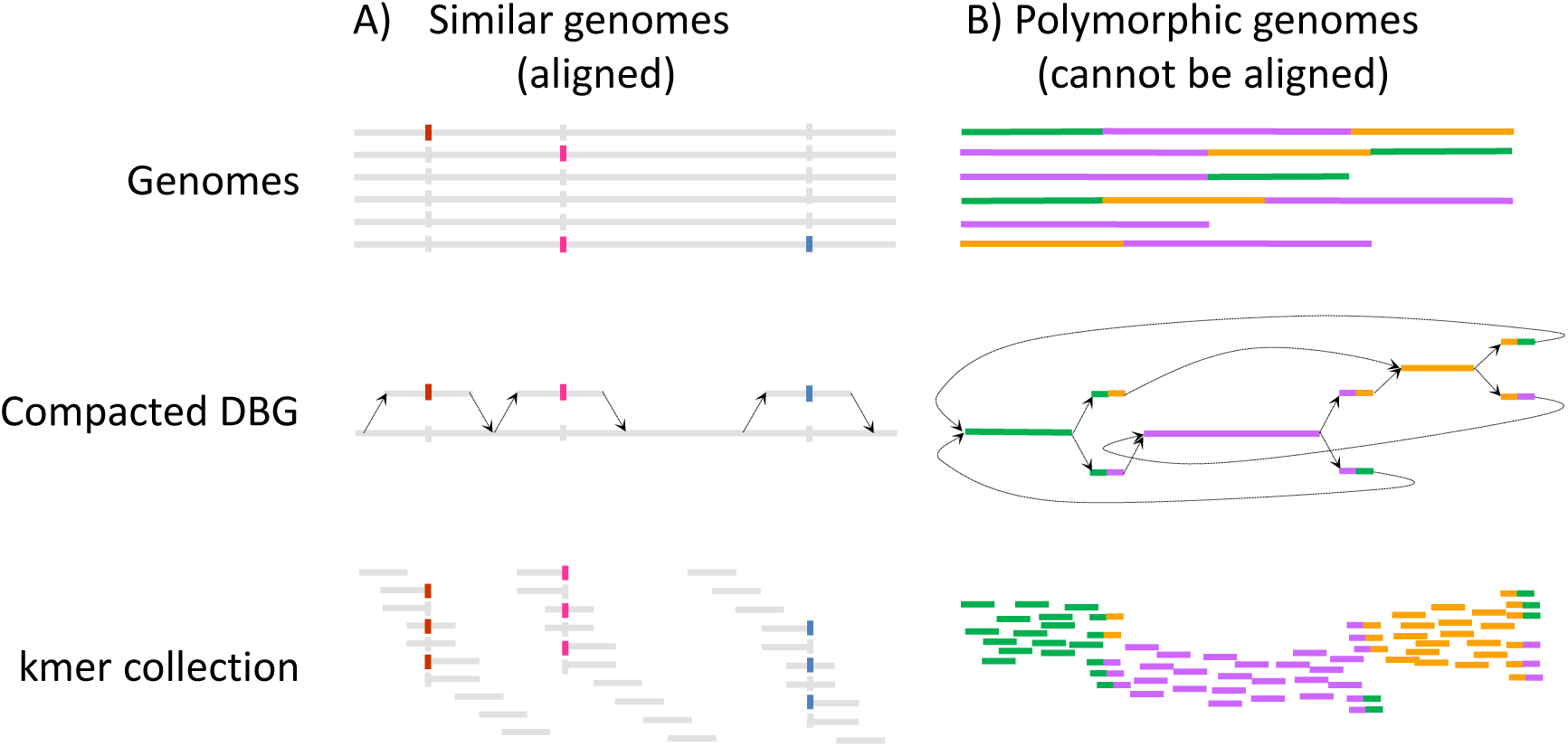
Alignment to a reference (when possible), DBG and kmers obtained for similar (A) and very polymorphic sequences (B). In the first case, the 3 loci represented as polymorphic in the alignment lead to 3 bubble patterns in the DBG, and numerous redundant kmers. In the second case, genomes are so polymorphic that an alignment is not possible. The DBG summarizes well the common regions and the links between them, while the collection of unique kmers still contains redundancy.

As variability across individual genomes increases and alignment of the genomes becomes ill-defined (Figure 2B), the DBG drifts away from a path to accommodate local variation beyond isolated SNPs. Fixed-length kmers are also able to represent this variation but still contain a lot of redundancy: all kmers with a given color arise from the same larger colored segment (or junction between segments). They correspond to the same unitig, and their presence/absence across strains is the same. By contrast, the DBG exploits the fact that some regions can be more or less polymorphic across genomes to compact redundant kmers into single longer non-redundant unitigs: their presence/absence across strains is different – unless the corresponding regions are present in the exact same set of genomes because of linkage disequilibrium (LD). In the extreme case where genomes in the panel have so little in common that no compaction is possible in the DBG, the unitig representation reduces to the fixed-length kmer representation.

In this sense, unitigs always represent the best of both worlds between a SNP-based representation of genetic variation and one based on a set of unstructured fixed-length kmers. It results in a locally optimal resolution: regions of the genome which are conserved across individuals are represented as single long words while regions which are too variable are fractioned into shorter structured kmers.

In addition to removing redundancy compared to fixed-length kmers, DBGs maintain an information regarding how kmers follow each others in the panel, and can be used to interpret those whose presence in the genome is associated with resistance by visualizing the proportion of resistant strains in which they are present. We use these facts in Section 2.5.2 to interpret our results.

#### 2.1.2 Choice of k

Each choice of a fixed-length k leads to a different DBG (Supplementary Figures 1), and there is no general rule as to how to choose k. Small values of k produce very connected sets of non-specific kmers which fail to represent the specificities of the data. In particular, any region larger than k which is repeated in two different parts of the genome creates a cycle in the DBG. On the contrary, large values of k can fail to create 2 different nodes for 2 different SNPs separated by less than k bases. In this case, the 2 SNPs will be considered as a unique variant. We tested a few values of k and judged by the general aspect of the DBG obtained on our panel and by the GWAS performance, as detailed in Supplementary Figures 1 and 2. We fix k to 31 for the rest of this study, as this value leads to both an exploitable topology for the DBG built on the gyrA gene, and good performances on GWAS. We found our results to be robust to small variations of k. We discuss the effect of k in more detail in Section 3.1.

### 2.2 Testing procedure

We build our test using a linear model relating resistance phenotypes to a candidate genetic determinant and population structure. Let *n* be the number of observed samples (*i.e*., strains with available genome and phenotype). When testing any particular haplotype (presence/absence of a unitig or a fixed-length kmer in the genome) for association with the resistance phenotype, we use the following model:

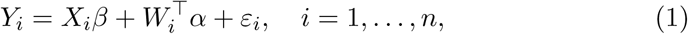

with 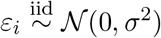, *σ^2^ >* 0. For any sample *i*, *Y_i_* is a binarized antibiotic susceptibility status: 0 for susceptible strains and 1 for non-susceptible (resistant and intermediary) strains, *X_i_* is 1 when the sample has the major version of the haplotype, 0 otherwise. We discuss the set of tested candidates *X* in Section 2.3. *β* is the effect of the haplotype on the phenotype, *W_i_* ∈ ℝ^*l*^ is a factor representing the population structure, *α* ∈ ℝ^*l*^ is the effect of this population structure on the phenotype. We choose to use a linear model rather than a logistic one even though our outcomes *Y_i_* are binary: we tried a logistic model in preliminary experiment, but obtained worse detection performances. Many combinations of *X* and *W* factors indeed led to poorly conditioned optimization problems and poor numerical solutions. The logistic model also led to much longer computation for the test.

Our objective is to detect haplotypes whose presence in the genome is associated with antimicrobial resistance. Formally for each haplotype *X*, we test *H_0_*: *β* = 0 versus H_1_: *β* ≠ 0 in model (1).

It is well known from the human GWAS literature that spurious associations can be detected if the effect of the population structure is not taken into account (Balding, 2006; Zhou and Stephens, 2014; Widmer *et al*., 2014). For example, assume a clade contains only resistant individuals because a mutation acquired by a common ancestor of this clade confers resistance. Then all other mutations which are acquired later in evolution and are more present in the clade will also be found to be associated with the resistance phenotype. Population structure can be very strong within bacterial strains (Earle *et al*., 2016; Falush and Bowden, 2006). We estimate this structure from the whole design matrix **X** ∈ ℝ^*n*×*p*^, where *p* is the number of unitigs or kmers (as discussed in Section 2.3, **X** typically has several identical columns). We evaluated with three models on both simulated and real data: (i) no correction, (ii) fixed effect α and (iii) random effect α. Denoting **X** = *U* Λ *V^T^* the singular value decomposition (SVD) of **X**, we use *W* = *U_q_* (the matrix formed by the first *q* columns of *U*) in the fixed effect model and 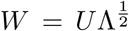 in the random effect model. For the first two models, we compute p-values for *H_0_* using a likelihood ratio test. For the random effect model, we use the bugwas implementation of Earle *et al.* (2016) to test *H*_0_, providing a pre-computed population structure *W*. Note that bugwas also offers to detect “lineage effect”, namely columns of the population matrix *W* which are associated with resistance, as a mean to avoid throwing away candidates whose association is explained away by the population structure: some of them could actually be causal, and bugwas would return the whole lineage along with correlated candidates as a lower resolution entity.

### 2.3 Genome-wide variant matrix building

One goal of this paper is to illustrate the advantage of testing unitigs rather than fixed-length kmers for association with antimicrobial resistance. To do so, we need to represent both fixed-length kmers and unitigs as 2-levels factors coding for the presence/absence of kmers/unitigs in model (1). More precisely, we consider both as generalized haplotypes with two alleles: presence or absence of the kmer or unitig in the genome. We then express each haplotype as a binary vector *X* ∈ {0,1}^*n*^, with *X_i_* = 1 if sample *i* has the minor allele (the less frequent one across the dataset), 0 otherwise. Consistently with Earle *et al.* (2016), we refer to such a binary vector as a pattern in the remainder of this paper, to emphasize the difference with the actual kmer or unitig they represent. Different kmers or unitigs can indeed be represented by the same binary vector because their presence/absence pattern across the genomes is the same. We only perform one test for each unique pattern (presence/absence binary vector), but retain the link between each pattern and the corresponding kmers and unitigs for later interpretation.

Both fixed-length kmers and unitigs lead to the same set of distinct patterns (represented by vectors in {0,1}^*n*^) across the genomes. Indeed, every unitig represents (at least) one fixed-length kmer, and conversely every fixed-length kmer is represented by one (single) unitig. As a consequence, the set of patterns tested for association with microbial resistance is identical for the two representations, which further illustrates the fact that using unitigs does not remove information compared to fixed-length kmers.

Every pattern we test often corresponds to a large number of fixed-length kmers. Many of them can come from a single longer sequence of DNA which is either entirely present or absent in each genome of the panel: in this case, they all map to the same unitig. This redundancy is a nuisance because it amounts to artificially fractioning a single pattern into several pieces only because we are not working at the right resolution. For example, the SNP on Figure 1 can be represented by one long kmer (unitig) whose only variation across all genomes is at the position of this SNP. Likewise, a unitig can correspond to a gene which is present in some of the genomes but not all of them. In both cases, breaking the unitig into several shorter fixed-length kmers does not bring any additional information and makes the results harder to interpret.

Each pattern in turn typically corresponds to a much lower number of unitigs than kmers. By construction, two unitigs related to the same pattern cannot correspond to overlapping words – they would have been compacted as a single longer unitig otherwise. They are only redundant in the sense that a genome contains one of the unitigs if and only if it contains the other. This redundancy can be dealt with by inspecting the DBG, as we discuss in Section 2.5.2. The reasons can range from nearby nodes being separated by a rare variant, to two separate genomic regions being in LD.

We build a single compacted DBG from 282 *P. aeruginosa* genome assemblies (see section 2.4) using the kissplice software, version 2.3.1 (Sacomoto *et al*., 2012). A specific aspect of our approach is that we build our compacted DBG from assembled genomes (more precisely, from contigs) rather than from primary sequence reads. This allows us to avoid dealing with sequencing errors, which are present in reads but are mostly eliminated during the assembly process. We choose kissplice settings in order to have no filter on the kmer frequencies or occurrences (−c 0 −C 0.001) and build one DBG per tested kmer length: k=13, 15, 17, 19, 21, 31, 41, 51 and 61 pbs. All resulting fixed-length kmers and DBG unitig sequences are then mapped without mismatch to the original genome assemblies using Bowtie 2 (Langmead and Salzberg, 2012) in order to determine the presence or absence of each kmer and unitig in each genome, as this information is not provided by kissplice.

### 2.4 Dataset

We use a panel of 282 strains of *P. aeruginosa* species, a ubiquitous bacterial species responsible of various infections, highly adaptable thanks to its ability to exchange genetic material. The species accessory genome is particularly important, in terms of size and diversity, and carries a large part of the genetic determinants already described to confer resistance to antimicrobial drugs (Jaillard *et al*., 2017). This strain panel was gathered from two collections including mostly clinical strains: the bioMérieux collection (*n*=219) (van Belkum *et al*., 2015) and the Pirnay collection (*n*=63) (Pirnay *et al*., 2009). Genomes were sequenced on Illumina HiSeq 2500, assembled using a modified version of the IDBA_UD assembler (Peng *et al*., 2012), and annotated for the identification of core and accessory genes (van Belkum *et al*., 2015). Both sequencing and assembly are available on NCBI with accession number PRJNA297679.

Antibiotic resistance phenotypes were obtained by broth dilution assays complemented with VITEK2 testing (bioMérieux, Marcy-l’Étoile, France), for several drugs commonly used in *P. aeruginosa* infections, including amikacin (280/282 strains) and levofloxacin (117/282 strains) (van Belkum *et al*., 2015). A minimal inhibitory concentration (MIC) value was thus available for all the characterized strain/antibiotic couples. Clinical and Laboratory Standards Institute (CLSI) guidelines were applied on the resistance data to determine susceptibility or non-susceptibility (phenotypic data in Supplementary Table 1). The reader is referred to van Belkum *et al.* (2015) and Jaillard *et al.* (2017) for more information on all strains and their analysis.

### 2.5 Evaluation

We evaluate two complementary aspects of our unitigs. First, we verify that when used in GWAS, they lead to the detection of true genetic determinants on both simulated and real data, under different population structures. Then we assess how insightful the representation is and what type of event underlies each tested pattern.

#### 2.5.1 Ability to detect variants associated with resistance

Quantifying how well a detection method works is difficult, as not all genetic determinants of antimicrobial resistance are known. If a method calls an association between resistance and a particular variant which was never described as causal it may be a false positive but it may also be because the method discovered a new unreported mechanism. We therefore choose to evaluate how well our test detects true determinants on three complementary indicators.

First, we simulate resistance phenotypes based on our real genomes, arbitrarily fixing which patterns *X* built in Section 2.3 have a non-zero effect on the phenotype *Y*. Let **X̃** be an *n* x *q* matrix whose columns are the unique patterns (in contrast with **X** ∈ ℝ^*n*×*p*^ whose columns correspond to typically non-unique kmers or unitigs), *β* is an ℝ*^q^* vector of the corresponding effects. We sample the phenotype *Y_i_* of each sample *i* from a multivariate logistic model:

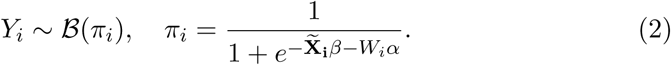

Using this set of positive and negatives, we can plot a Receiver Operating Characteristic (ROC) curve for each of the three methods introduced in Section 2.2. The simulation is multivariate, accounting for the fact that resistance can stem from a combination of causes rather than a single one whereas we are using univariate model (1) for our test. Since we use a logistic model which is the generalized linear model (GLM) of choice to handle binary outcomes, as opposed to the linear model which we use for convenience in our test, it also takes into account the potential misspecification between the model underlying our procedure and the actual distribution of the data. On the other hand, the conclusions we draw from this simulation are contingent upon the capacity of the logistic model (2) to represent the relationship between haplotype and phenotype.

Using the true phenotype data for both amikacin and levofloxacin resistance, we also evaluate a metric based on libraries of known genetic determinants of resistance (Jaillard *et al*., 2013) (mentionned thereafter as reported causal variants) which we use as our positive set. In this case we do not need the assumptions made in the simulation, but we lose the exact knowledge of which haplotypes are negative, *i.e.,* have no effect on the phenotype: some selected patterns may not be linked to any know genetic determinant of resistance just because there are still unreported. Instead of ROC curves, we therefore resort to plotting the true positive rate (TPR) – using identified and hence known positives – as a function of the number of positives called by the method – the false positive rate corresponding to this number being unknown. Assigning each selected pattern (which can represent several mutations or presence of accessory genes) to a true or false status requires a mapping step and some type of approximation: we choose to identify a pattern as a true determinant if it corresponds to at least one kmer/unitig which maps to a known genetic determinant from a resistance gene sequence database (Jaillard *et al*., 2013).

Finally using the true phenotype data for amikacin resistance, we plot the proportion of reported causal variants recovered as a function of the number of kmers or unitigs called positive. We restrict ourselves to the million kmers (resp. unitigs) with the lowest p-values. While the first two metrics focus on unique patterns and do not distinguish between kmer and unitig encoding (both leading to the same set of patterns), this third metric allows us to compare the number of kmers and unitigs that need to be inspected to identify a given proportion of all reported causal variants. This number can be different as each presence/absence pattern corresponds to different numbers of kmers and unitigs.

#### 2.5.2 Making sense of the selected patterns using the compacted DBG

The analysis we describe in Section 2.5.1 is necessary because we need to verify that our test actually discriminates between patterns corresponding to causal variants and those not corresponding to any causal variant. It is however not sufficient to ensure that our procedure is suited to identifying unreported genetic determinants of antibiotics resistance: to be able to perform this analysis, we had to define which patterns were true determinants using annotated SNPs and genes known to be linked to resistance. In addition to being approximate, this definition cannot be used to go beyond recovering existing determinants.

In order to perform this task, we must be able to interpret the selected patterns. Assuming a pattern is found to be associated with resistance in our test, its interpretation in a fixed-length kmer paradigm can be cumbersome: it typically requires to map all kmers corresponding to the pattern to all genomes– as there is no single reference genome in this context – and to make sense of these mappings. For example, one may find that several of these kmers map to similar regions or annotated genes in all genomes. The task can be heavy as each pattern is typically associated to a large number of redundant fixed-length kmers.

Annotation of our unitigs is easier for three reasons. First, the number of unitig sequences to be mapped is much lower than the number of fixed-length kmers, as illustrated on Figure 3. Second, unitigs are longer than kmers, making them more likely to map to a unique region in the genome. Finally, the DBG itself and its colored version (Iqbal *et al*., 2012) can help us understand which type of event is associated with a unitig. The colored DBGs we use rely on node sizes to represent allele frequencies, *i.e.,* the proportion of genomes containing the sequence. They also rely on node colors to represent the proportion of resistant strains containing the corresponding unitig, countinuously interpolating between a red node for unitigs found in resistant strains only and a blue node for those found in susceptile strains only.

**Figure 3:**
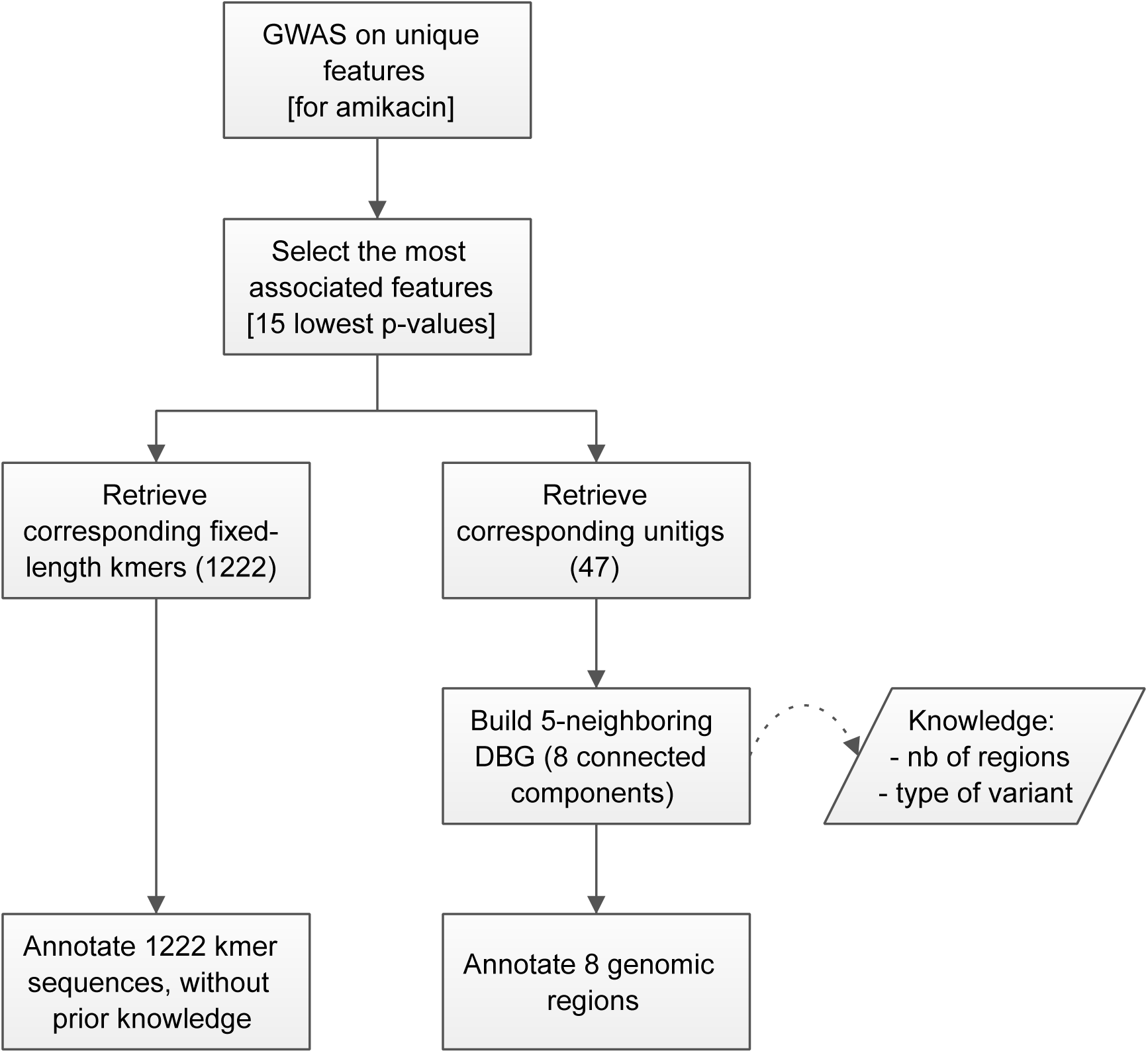
Flowchart of post-processing. The flowchart is illustrated with the results obtained for amikacin resistance: settings are given between brackets while resulting numbers are given between parenthesis. The annotation burden is lighter when using the DBG unitigs than fixed-length kmers. Indeed in the case of amikacin resistance, the number of kmer sequences to map against all genome exceeds 1000, while using the DBG unitigs we map no more than 47 unitig sequences and can also rely on the identified 8 genomic regions for a complementary interpretation.

Concretely, we select a few patterns with lowest p-values from the GWAS results. We then retrieve all unitigs corresponding to these low p-value patterns–some unitigs can share their presence/absence profiles because of LD, and thus are duplicated. We build the subgraph of our colored DBG induced by these top unitig plus all their neighboring unitigs for a given neighboring size s. We refer to this subgraph as the s-neighboring DBG. This representation offers several advantages:

1. It can be done regardless of the association of the pattern with the resistance phenotype and whether or not any annotation is available for the studied genomes. Its topology reflects the nature of the variant: bubbles for example correspond to SNPs while paths represent gene insertion.
2. Node colors visually help understand which unitigs or more complex subgraphs are associated with resistance. This allows us to identify bubbles (*e.g*. SNPs or indel) whose branches differentiate phenotype status, and can still be done when no genome annotation is available, using only the strain phenotypes.
3. Top unitigs which are close to each others in the genomes will be gathered into connected components of the induced subgraph. These components may represent well-defined genomic regions such as genes or mobile genetic elements – not all connected components will correspond to genomic regions however: some may result from repeated regions in the genome. On the other hand, unitigs mapping to different connected components – distant neighborhoods – carry information on LD, *i.e.,* separate haplotypes which happen to be present in the same set of samples, possibly because of the population structure.

## 3 Results

We describe the results obtained in our experiments on simulated and real antibiotic resistance phenotypes. We study both the ability of the unitigs to detect causal variants when used in GWAS and the interpretability of the detected objects.

### 3.1 Extracting fixed-length kmers and unitigs from complete genomes

The length k of the kmers used to build the DBG determines how the DBG represents our set of genomes and its ability to provide some level of compression. Small values (below 20) generate words of low complexity which are highly repeated in the genomes, creating numerous loops in the DBG. Consequently, the graph is hardly compacted, as it is very connected and contains few linear parts. For k=15, we only count twice more kmers than unitigs (34 M versus 15 M). As k increases, the number of kmers increases but they become more specific and less repeated within genomes, leading to better levels of graph compaction. For k=41, we obtain 62.5 M kmers and 2.2 M unitigs. More generally, panel A of Figure 4 shows that as k increases, the number of kmers increases whereas the number of unitigs remains stable.

**Figure 4:**
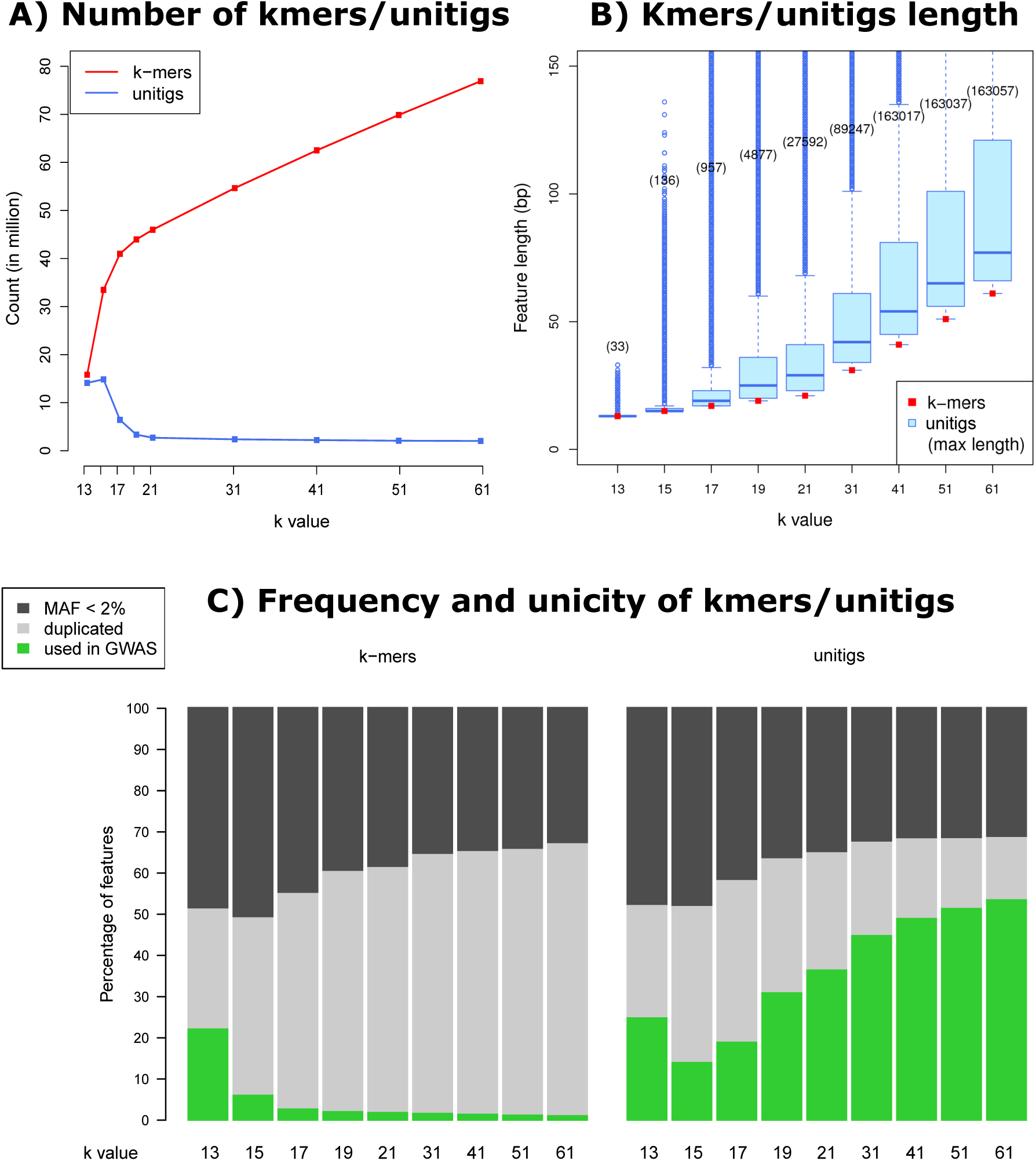
Preprocessing. Panel A shows the number of fixed-length kmers (red) and unitigs (blue) in the data as a function of *k*. Panel B shows the corresponding distribution of variable length kmers associated with each unitig. Panel C shows which proportion of kmers and unitigs correspond to unique presence/absence patterns in the data.

Simultaneously, panel B of Figure 4 shows that increasing k leads to unitigs of increasingly variable size - larger or equal to k by construction. For k=41, the median length of unitigs is 54 and the longest unitig is 163017 bp long. This illustrates both the redundancy of the fixed-length kmer representation and the capacity of unitigs to produce descriptors whose resolution is adapted to the local variation observed across the genomes.

Panel C of Figure 4 represents for each k the percentage of kmers or unitigs which we filter out from our GWAS because their minor allele frequency (MAF) is too low (dark grey). Furthermore as discussed in Section 2.3 several kmers or unitigs can have the same presence/absence pattern on a given set of genomes, so we also represent the proportion of kmers or unitigs which are filtered out from our GWAS because they correspond to duplicated kmers or unitigs (light grey). As expected, this proportion is much larger for fixed-length kmers than for unitigs: a large fraction of fixed-length kmers associated with a single pattern are summarized as a single unitig. This is consistent with the observation that the number of fixed-length kmers is much larger than the number of unitigs but that both representations ultimately lead to the same number of unique patterns.

### 3.2 Phenotype simulation study

We generate synthetic data with two scenarios under model (2) to illustrate the capacity of our test to detect patterns associated with resistance and the importance of adjusting for population structure. This will also help interpret results on real data in Section 3.3.

We use the design matrix **X** built from our panel of *P. aeruginosa* genomes. We compute its singular value decomposition **X** = *U* Λ *V^T^* and set 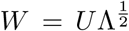.

Our first scenario is intended to illustrate the case where there is a population effect on the observed resistance (some clades are enriched or depleted in resistant samples) which is not explained by the set of patterns in the tested design *X*. In practice, this could be a non-genetic (*e.g*. environmental or batch) effect. More importantly, this could happen if some genetic determinants are not included in the model used for testing. This is likely to be the case when we use model (1) which is univariate, *i.e*., which only considers one pattern at a time. For example, it could be the case that one mutation *A* causing resistance was acquired by the ancestor of a clade and transmitted to its descendants: the clade would then be enriched in resistant individuals. If a second mutation *B* not related to resistance is acquired by a close descendant of the common ancestor and transmitted, many samples from the clade will also have mutation *B*. A univariate test of association of *B* with resistance will not account for *A*. If the test does not account for population structure either, it may assign a smaller p-value to *B* than to other mutations with an actual causal effect, *e.g.* because these mutations involve fewer individuals, which leads to a lower power to detect true determinants.

To simulate this scenario, we arbitrarily assign two columns of *W* (the second and the sixth) to have non-zero effects *α*, so *l* = 2. By construction, the first columns of W represent a large fraction of the variation across strains. A non-zero effect *α* in the GLM (2) used to simulate resistance phenotypes therefore makes resistance associated with the population structure. We then select 10 distinct patterns from **X̃** as true determinants (*i.e*., coordinates *j* ∈ 1,…, *r* associated to non-zero effects *β_j_*). To do so, we compute the largest dot product of each pattern with the first six columns of *W* (two of which have non-zero effects *α*), and choose our true determinants among those whose largest dot product is below the fifth percentile of dot products calculated across all patterns. This allows us to simulate the case where true determinants are independent from the population structure (their effect is not inflated by the *W α* term). The odd ratios *e^β_j_^* are fixed to 6 for these patterns. We also randomly select 290 patterns from **X̃** as non-determinants, *i.e.,* with a *β_j_* = 0 effect in the model, so *r* = 300 in our simulation. The population structure can lead to spurious discoveries, as we do not control the dot product between columns of *W* and these patterns with zero effect. Finally in order to control the amplitude of the population effect, we normalize *W α* to 6 times the median value of the |**X̃^j^***β_j_*| across non-zero *β_j_*, where **X̃^j^** denotes the *j-*th columns of **X̃**

We then apply the three versions of our univariate test described in Section 2.2 to each of the patterns. For the fixed effect correction, we use the first 10 columns of *W*, and for the random effect correction we provide the entire *W* matrix to bugwas. We perform 100 repetitions of this simulation, and plot a Receiver Operating Characteristic (ROC) curve after pooling the results (Supplementary Figure 3). As expected, the test which does not account for the population structure has very low power to detect patterns associated with the phenotype: by construction, some patterns with zero actual effect have large dot products with *W α* which inflates the estimate of their effect and leads to false discoveries. Taking the population structure into account in the model improves the power by limiting this inflation.

Our second scenario is meant to illustrate the case where there is little population effect observed on the phenotype except for that caused by the association of modeled causal patterns *X* with *W*, i.e., outside of **X̃***β* in (2). In other words, we assume that all the imbalance in proportion of resistant samples across clades is explained by patterns in the design **X̃**. This can happen if most of the true causal patterns are not too related to the population structure, *e.g*. because they appeared by homoplasy on several unrelated individuals and there is no imbalance. In this case, correcting for the population structure can decrease the estimated effect of causal patterns which do have some association with this structure, *i.e.*, which were acquired by ancestors. To simulate this scenario, we use the same setting as before but we select the 10 true determinants among those that have a large dot product with W rather than a small one, and set all *α* effects to zero. We apply the same three tests as in the previous scenario over 100 data generations and plot a ROC curve. This time, we observe the opposite effect as in the previous scenario: correcting for the population structure decreases the power to detect true determinants. Assuming there is a population effect when there is no such effect in reality leads to artificially deflating the estimated effects of patterns which are associated with the population structure.

### 3.3 Application on real data

We then turn to results obtained from real actual amikacin and levofloxacin resistance measured on this panel.

#### 3.3.1 True positive rate vs number of positive predicted

Supplementary Figure 4A is produced by bugwas, and shows the p-value of the test for association of each column of *W* with the phenotype (Earle *et al*., 2016). In the case of amikacin, two columns are found to have a significant effect at level 0.01, whereas all columns have p-values larger than 0.01 in the case of levofloxacin. Accordingly, Supplementary Figure 4B shows that correcting for population structure increases the proportion of known genetic determinants of resistance to amikacin recovered for every number of predicted positives, but decreases this proportion in the case of levofloxacin.

The results on the amikacin resistance phenotype are consistent with our first simulation, where the population structure had a non-zero effect *α* on resistance: the estimated effect *β̂* of true determinants which are not associated with the population structure (low dot product between *X* and *W α*) is unaffected by the presence of a population effect while the *β̂* of some patterns confounded with *W α* but with zero actual effect *β* are inflated. Consequently, the true determinants are not ranked among the first patterns, leading to decreased performances on Supplementary Figure 4. Correcting for the population structure limits this inflation of *β̂* for negative patterns associated with the population structure.

Conversely, assuming there is indeed no unmodeled effect of the population structure on levofloxacin resistance, corrected models may just underestimate the effect of true determinants whose presence is associated with the population structure, as in our second simulation. For example, if a causal SNP is shared by a clade which is consequently enriched in resistant samples and all the other SNPs shared by this clade also are causal, correcting for the population structure only decreases the estimated effect of the true determinant, leading to decreased performances on Supplementary Figure 4.

The random effect approach of bugwas is a good choice on both simulated (Supplementary Figure 3) and real data (Supplementary Figure 4B) regardless of the effect of the population structure on the phenotype: it outperforms both the uncorrected and the fixed effect approaches in the presence of a population effect and is only moderately affected by the absence of such effect.

#### 3.3.2 True positive rate vs number of explored features

The analysis of Section 3.2 and 3.3.1 establishes that representing genomes by their unitig content in GWAS allows to discriminate between (reported) causal variants and other variants (including non-causal and unreported causal variants in our experiment on real resistance phenotypes). However necessary, this result does not illustrate an advantage of unitigs compared to fixed-length kmers as both lead to the same set of presence/absence patterns and the analyses of both Sections only involve these patterns.

By contrast, Figure 5 shows the TPR for detecting reported causal variants for amikacin resistance as a function of the number of kmers and unitigs called positive – from 1 to 10^6^. In other words, this metric indicates which proportion of reported causal variants is recovered after inspecting a given number of elements. The unitigs perform much better than the kmers in this metric because every false positive pattern typically leads to a very large number of false positive kmers, and a lower number of false positive unitigs. This illustrates the fact that manipulating kmers is more cumbersome than unitigs as it is necessary to inspect, map and annotate more kmers than unitigs to recover the same number of causal variants.

**Figure 5:**
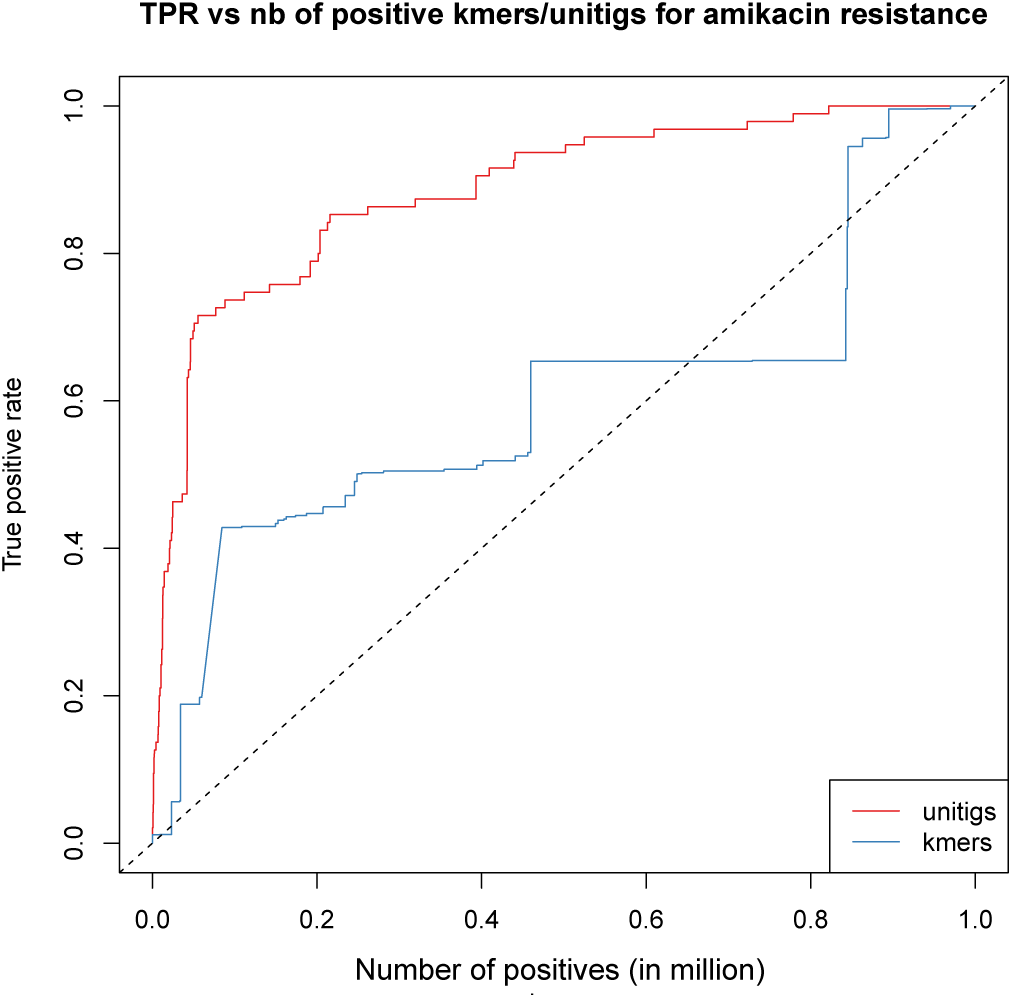
Proportion of true positive versus number of predicted positive kmers and unitigs for the first 10^6^ positive calls using bugwas on the amikacin resistance phenotype.

#### 3.3.3 Analysis of the selected haplotypes

We build the 5-neighboring DBGs from the 15 patterns with lowest bugwas p-values, for both amikacin and levofloxacin resistance, as described in Section 2.5.2. Using the uncorrected or fixed effect approaches leads to very similar lists of 15 patterns.

The top 15 patterns correspond to 47 unitigs for amikacin (resp. 22 for levofloxacin) or 1222 (resp. 262) kmers. The 5-neighboring DBG induced by these unitigs has 8 (resp. 6) connected components whose unitigs consistently map to a small number of annotated events (Supplementary Figures 5 and 6, Supplementary Tables 2 and 3).

The annotation of the 8 components found for amikacin highlights the importance of the accessory genome in resistance. Indeed, all top patterns map within or near mobile elements: more than half the connected components represent coding or non-coding neighborhood of transposase or integrase. By contrast, half of the 6 components found in the levofloxacin experiment represent SNPs in core gene, recognizable by paths of node with a high prevalence in all strains (violet nodes), and forks that split between a red (resistant phenotype) and blue (sensitive phenotype) path. This matches the current knowledge about levofloxacin resistance mechanism, mainly based on target alteration.

As discussed in Section 2.5.2, the few connected components induced by the top 15 patterns are much easier to interpret than the corresponding large sets of fixed-length kmers. We select 6 of these connected components (Supplementary Figure 5g, a, h and 6b, c, f) and extend their neighborhood up to distance 20 rather than 5 to illustrate the large variety of variants which are selected by our procedure.

##### SNP in an accessory gene (amikacin)

Figure 6A contains a quasi linear structure which evokes a polymorphic gene. The purple color of the structure suggests that the gene is more present in resistant than in sensitive samples, but that the differential of presence is not very important – the nodes would be red otherwise. In the middle of this structure (green box on the figure), the path forks into one blue and one red node, which suggests we have identified a SNP whose presence is associated with amikacin resistance. Note that we are able to make this interpretation regardless of any gene annotation, just by analyzing the topology of the graph component enriched by strain resistance information. Mapping the unitig sequences of this component onto our annotation reveals that the subgraph corresponds to the AAC accessory gene, whose presence is indeed known to be involved in *P. aeruginosa* resistance to amikacin. However, the selected event here is not the presence of the gene but the particular SNP within this gene.

**Figure 6:**
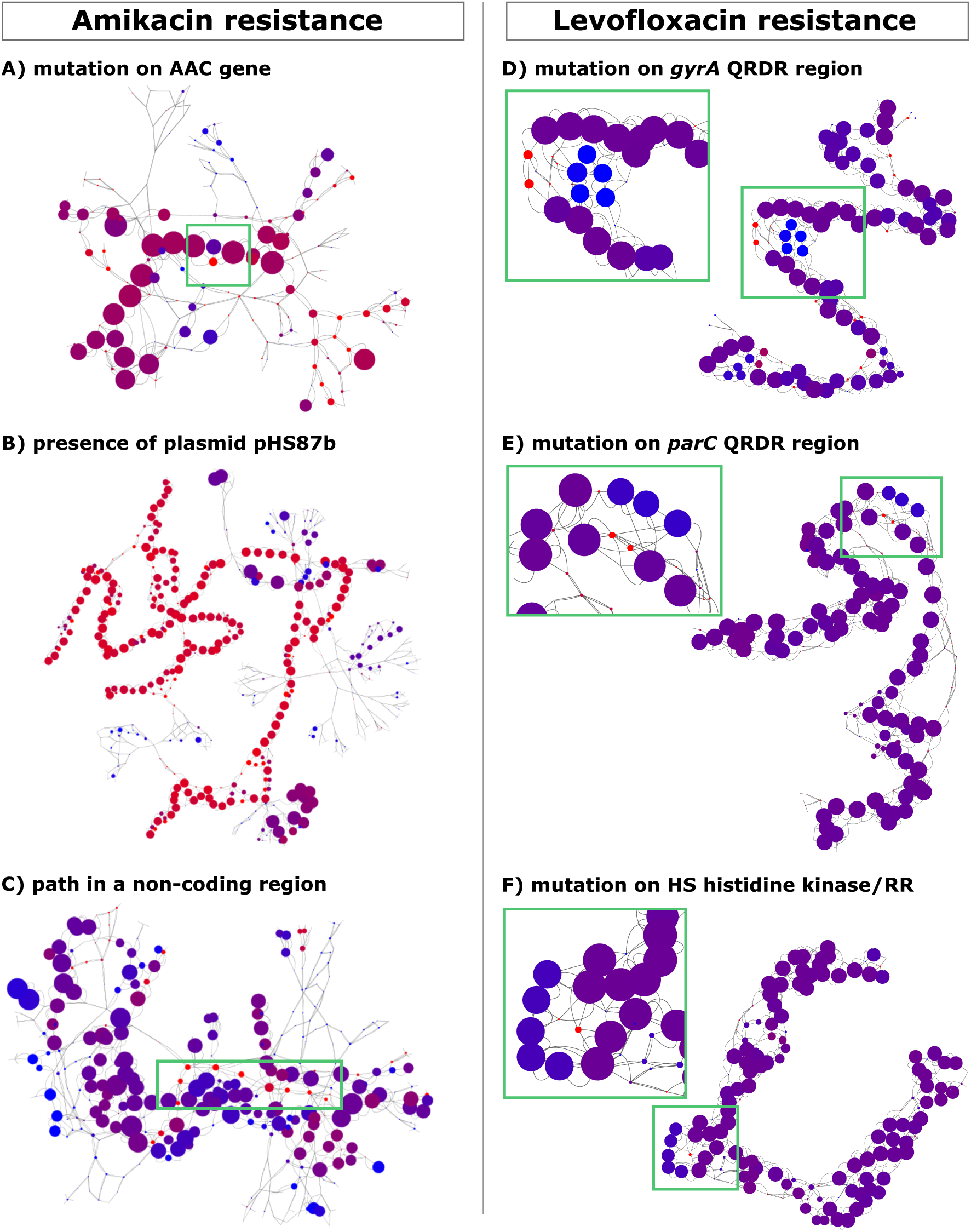
Neighboring De Bruijn subgraphs. Subgraphs of the De Bruijn graph obtained by retaining nodes separated by less than 5 edges from a node corresponding to a pattern whose p-value for association with resistance is among the 15 smallest. Node size represent the frequency of the corresponding haplotype, node color the proportion of resistant/sensitive ratio of samples containing this haplotype, from red for resistant only to blue for sensitive only. The left panel shows the result for amikacin resistance, the right panel for levofloxacin resistance.

##### SNP in a core gene (levofloxacin)

Components D to F of Figure 6 describe SNPs in core genes. Like in the previous AAC SNP example, each of these subgraphs is a linear structure in which most nodes are present in the same proportion of resistant and sensitive individuals. The linear structure contains a fork which separates resistant (red) and sensitive (blue) samples. Mapping the unitigs on sample genomes reveals that the first two components represent the well-known gyrA (D) and parC (E) quinolone resistance-determining region (QRDR). The third subgraph corresponds to a gene which is not present in our resistance database: the hybrid sensor histidine kinase/response regulator (HS histidine kinase/RR). This gene may be found associated with resistance to levofloxacin because it is in LD with a causal region, or may be itself causal.

##### Whole plasmid (amikacin)

Figure 6B shows a connected component with mostly red nodes assembled in a linear structure suggesting that this entire structure, as opposed to a point mutation, is involved in the detected event. This is in clear contrast with Figure 6A, where most of the linear structure is purple with a localized fork involving one red and one blue node. The unitigs of this subgraph corresponding to the top 15 patterns map to the pHS87b plasmid, which was recently described as being involved in resistance (Bi *et al*., 2016). Our representation extracts the whole plasmid, with both its coding and noncoding regions which makes it easier to understand that the selected patterns correspond to an integration of this plasmid.

##### Noncoding region (amikacin)

The unitigs of the component represented in Figure 6C map to a noncoding region in the *P. aeruginosa* genomes. Interestingly, this region contains a path of unitigs strongly associated with resistance (colored in red). Not all of these unitigs belong to the top 15, but the DBG view highlights this long linear structure. This haplotype is not compacted as a single unitig because it is not either present or absent in each genome: some only contain parts of this haplotype.

##### Alternative approaches

Our approach is able to select and detect any kind of event where current methods could be limited to some regions or patterns. SNPs called against a reference genome are of limited interest in the context of *P. aeruginosa* because of the size of the species accessory genome; causal variants in the accessory genome not represented by the chosen reference would not be detected at all. Gene presence/absence and SNPs called in the pangenome would miss all events in noncoding regions, by construction. Even assuming that only coding regions are causal, the noncoding region may have a strong association with resistance because of LD, and be among the top patterns in our test whereas the coding region is not because of noise, finite sample or model misspecification. Methods targeting only coding regions would miss the marker in this case. Finally, the gene presence/absence approach would miss the SNP that we identify in the AAC accessory gene. It could have detected the presence of the full gene as being associated with resistance to amikacin, but with less power: only one mutated version of the gene is involved in resistance. Fixed-length kmer approaches are able to target any region of all genomes. However in the case of an event defined by the presence of a complete plasmid such as pHS87b, a fixed length kmer representation would lead to identifying disconnected regions. Identifying the whole plasmid rather than sets of disconnected hits makes it easier to understand which mechanism underlies the selected patterns.

## 4 Discussion

We have introduced unitigs as a new and efficient mean to represent candidate genetic determinants in GWAS. Unitigs correspond to variable length kmers: genomic regions which are constant across samples map to single long kmers while more polymorphic regions are supported by several shorter kmers, leading to higher resolution. This representation generalizes both SNPs obtained by alignments against reference genomes and fixed-length kmers. Compared to the former, it is more flexible and can deal with highly plastic genomes. Compared to the latter, it is less redundant and leads to a drastic reduction in the number of candidate entities that need to be tackled without loss of information, leading to easier computation and interpretation of the result. Furthermore, extracting neighboring De Bruijn subgraphs provides additional insight as to what type of genomic event underlies a unitig which is detected as being associated with a phenotype of interest. Experiments on *P. aeruginosa* illustrate that our representation is able to capture very different genomic features ranging from SNPs to large gene insertions.

We conjecture that using unitigs rather than fixed-length kmers could also yield better estimates of the population structure. Typical estimators of this structure are based on representations of the genomes by their haplotypes rather than their unique patterns to avoid down-weighting haplotypes which map to the same presence/absence profile. While duplicated unitigs only represent biological duplicates, *i.e*., regions in perfect LD, duplicates within kmers also account for neighbor sequence overlaps and can lead to arbitrary inflation of the weight of single long haplotypes. Validating our conjecture that DBG nodes provide better population structure estimates than kmers and lead in turn to more power for detecting genetic determinants requires simulation of synthetic genomes from a given phylogeny and will be the subject of future work.

Finally an important improvement would be to generalize our representation to paths or more general subgraphs of the DBG, *i.e*., to larger haplotypes defined by conjunctions of those represented in unitigs. This could help filter out minor variations in the genome which are unrelated to resistance but prevent long haplotypes to be merged into a single node. The De Bruijn neigbhoring subgraphs we selected in our experiments suggest that this configuration happens frequently in practice.

## Acknowledgements

The authors thank Sarah Earle, Chieh-Hsi Wu and Daniel Wilson for their insightful comments.

## Funding

LJ is funded by the ANR (MACARON project under grant number ANR-14- CE23-0003-01). LL is funded by the Brazilian Ministry of Science, Technology and Innovation (MCTI), under the Science Without Borders (CSF) scholarship grant process number 203362/2014-4. VL is funded by the Agence Nationale de la Recherche ANR-12-BS02-0008 (Colib’read).

